# Higher order spherical harmonics reconstruction of fetal diffusion MRI with intensity correction

**DOI:** 10.1101/297341

**Authors:** Maria Deprez, Anthony Price, Daan Christiaens, Georgia Lockwood Estrin, Lucilio Cordero-Grande, Jana Hutter, Alessandro Daducci, J-Donald Tournier, Mary Rutherford, Serena J. Counsell, Merixtell Bach Cuadra, Joseph V. Hajnal

**Author notes:** Corresponding author Email address (Maria Deprez).

## Abstract

We present a comprehensive method for reconstruction of fetal diffusion MRI signal using a higher order spherical harmonics representation, that includes motion, distortion and intensity correction. By applying constrained spherical deconvolution and whole brain tractography to reconstructed fetal diffusion MRI we are able to identify main WM tracts and anatomically plausible fiber crossings. The proposed methodology facilitates detailed investigation of developing brain connectivity and microstructure in-utero.

## 1. Introduction

Fetal diffusion magnetic resonance imaging (dMRI) can provide detailed insights into healthy development of brain connectivity and microstructure in the pre-natal period. Fetal diffusion imaging is however very challenging due to a number of issues: unpredictable fetal motion and maternal breathing; geometric distortion of echo planar imaging (EPI); spin history artefacts due to long relaxation times of water-rich fetal brain tissues; significant intensity inhomogeneity that modulates intesities of fetal brain MRI in an inconsistent manner in presence of motion; and limited spatial resolution and signal to noise ratio resulting from the small fetal head being embedded inside the mother’s body. Early studies therefore relied on sequences that required very short acquisition time [1] or selecting the datasets least affected by motion [2], thus greatly limiting spatial and angular resolution and number of fetal scans that were suitable for further processing. To address these challeges, methods for correction of motion and imaging artefacts in fetal dMRI have been developed.

### Motion correction and scattered data interpolation

The duration of acquisition of each single diffusion weighted EPI slice is typically around 200-300ms and most of the slices are therefore not significantly affected by motion. However, motion is almost always present between individual slices. Jiang *et al.* [3] therefore proposed to use slice-to-volume registration (SVR) to correct for between-slice motion, similar to previously proposed approaches for motion correction of structural fetal MRI [4, 5]. Unlike structural MRI, different dMRI slices correspond to different sensitation gradients which change with respect to fetal brain anatomy according to the motion parameters. Jiang *et al.* [3] therefore proposed to register the diffusion data to the reconstructed (non-sensitised) *b*_0_ volume (*b* = 0*s/mm*^2^) using normalised mutual information as a similarity measure, followed by a model-based approach, where each slice is registered to the simulated volume with corresponding sensitation gradient. Another challenge, reconstruction of diffusion tensors on an isotropic grid from data scattered in both spatial and angular domain, has been addressed by Jiang *et al.* [3] using an interpolation model and inverting a large sparse matrix to calculate a consistent isotropic volume of diffusion tensors. Oubel *et al.* [6] has shown, that co-registration of slices of arbitrary gradient sensitations using cross-correlation as a similarity measure is more robust than registering them to a *b*_0_ image using normalised mutual information and this approach is also successfully adopted in this paper. To perform scattered data interpolation, they perform radial basis function (Gaussian) interpolation in both spatial and angular domains. A model-based approach that combines interpolation using Gaussian processes in the angular domain with volumetric registration has been shown to allow for motion correction of adult data with very high b-values (up to *b* = 7000*s/mm*^2^) [7], however, fetal data are subject to much larger motion and due to limited signal to noise ratio maximum b-values for fetal dMRI are generally lower, more typically up to *b* = 1000*s/mm*^2^.

### Super-resolution reconstruction

Due to the insufficient signal-to-noise ratio and to improve coverage of the brain in the presence of motion, fetal dMRI has been traditionally acquired using thick-slice sequences. Super-resolution approaches have been proposed for fetal structural MRI [8, 9, 10, 11] and later also for fetal dMRI [12]. Though thick-slice approaches are still most common for acquisition of fetal dMRI in clinical settings, novel isotropic fetal dMRI sequences [13] are being developed as a part of developing Human Connectome Project^1^ (dHCP). However, even in this context, the concept of modeling of acquisition of the slice using a point-spread-function (PSF) based on principles of MR physics [10] remains valid. In this paper we therefore adapt such an inverse problem approach, which generalises the estimation of regular isotropic dMRI volume from scattered data to both anisotropic and isotropic acquisitions.

### Spatial and angular regularisation

In the case of super-resolution reconstruction of fetal structural MRI, the reconstructed isotropic volume needs to be regularised to prevent noise enhancement due to the inverse problem being ill-posed. Various regularisation terms have been proposed, including L2 norm [8], edge-preserving regularisation [9, 10] and L1 norm [11]. The most common regulariser in the angular domain is the Laplace-Bertrami operator [14]. In our current experiments we have not utilised regularisation and this will be investigated in the future.

### Geometric distortion

A major issue when reconstructing fetal dMRI is geometric distortion of the native images when acquired using EPI. We have addressed this problem by registration of acquired *b*_0_ images to reconstructed structural data that are not subject the distortion with a physically inspired Laplacian constraint imposed [15], and this approach is applied to diffusion sensitised volumes in this paper. Dynamic time-varying distortion correction approaches that promise further improvements are being developed as a part of dHCP [13, 16].

### Spin history effects

Motion of the fetal head relative to the scanner can result in uneven timing between successive excitations of the magnetization in any given brain location leading to spin history effects, which are exacerbated because T1 relaxation times of fetal brain tissues are significantly longer than for adult brain due to high water content of fetal brain tissues [17, 18, 19]. The sequence repetition time (TR) can be increased to reduce or avoid spin history effects, but this becomes impractical, as a lot of scanning time would be lost waiting for recovery of the magnetization, limiting the number of diffusion sensitization directions that can be acquired. Therefore a compromise has to be made between spin history artifacts, image quality and efficiency. This means that spin history correction has to be incorporated into the reconstruction of the diffusion volume. We have previously addressed this problem in structural MRI by a data-driven approach [10], where slice-wise multiplicative low frequency fields were estimated to match intensities of slices to the reconstructed volume. We have adopted a similar approach for dMRI in this paper. An alternative strategy is to model spin history effects based on estimated motion parameters and principles of MR physics, as we previously proposed for fetal functional MRI (fMRI) [19] and this remains an area of future research for fetal dMRI. When motion disrupts the acquisition to the point that individual slices are corrupted we employ robust statistics as proposed for fetal structural MRI in [10] and adapted for dMRI, to exclude the corrupted slices.

### Bias field

Acquisition is further corrupted by spatial inhomogeneity of receive properties (B1−) of the array coils placed around the maternal anatomy and, when operating at higher magnetic field strength, the transmit (B1+) field can become variable also. We employ the method previously proposed for fetal fMRI [19] and correct the combined B1 effects in the pre-processing step by estimating B1 field from *b*_0_ images and applying them to diffusion sensitised images. An alternative method has been developed by Seshamani *et al.* [20] and could be used instead if sufficient number of *b*_0_ stacks in different motion states are present. These approaches are based on assumption that the combined B1 is a multiplicative low-frequency field that is stationary in scanner space in the presence of fetal motion. However, it needs to be stated that B1+ effects are not simply multiplicative, potentially resulting also in contrast variation, thus these methods offer only an approximate solution.

### Beyond diffusion tensor fitting

Previous works for motion correction of fetal dMRI were limited to reconstruction of diffusion tensors [3, 6, 12], mainly due to the data quality not supporting higher order models. Diffusion tensors however do not allow for modelling of crossing fibers and are therefore too restrictive for state-of-the-art fetal dMRI acquisition sequences, such as the ones being developed for the dHCP. In adult studies, one of the most popular ways to parametrise dMRI in the angular domain is using Spherical Harmonics (SH) [14, 21], through non-parametric models such as Gaussian Processes [7, 22] have been used as well. It has been previously shown that fiber crossings can be identified in fetal dMRI by using a SH model [23]. However, this work relies on the assumption of no fetal motion in the data.

### Contribution of this paper

In this paper we propose a novel framework to perform reconstruction of motion-corrected fetal dMRI signal from scattered data using a spherical harmonics (SH) parametrisation of the diffusion signal. We show that given sufficient angular resolution and b-value, we are able identify anatomically plausible fiber crossings in fetal dMRI by performing constrained spherical harmonics deconvolution [21] of the reconstructed signal. We also show that intensity artefacts such as B1 inhomogeneity and motion induced spin history effects need to be corrected to obtain consistent reconstructions. Additionally, susceptibility-induced distortion correction also improves reconstruction of fetal dMRI.

## 2. Methods

### 2.1 Reconstruction of Spherical Harmonics representation of dMRI signal from scattered data

#### Spherical Harmonics representation

Our aim is to provide a robust methodology to flexibly support a wide range of diffusion analyses given data that is both scattered in space and may have irregular angular samples that differ from one spatial location to another. The use of a SH basis allows direct fitting to variable angular samples without the need for an initial interpolation step. We therefore seek to estimate a SH representation *C* = {*c_i,lm_*} of fetal dMRI signal 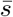 on a high-resolution isotropic spatial grid represented by index *i*

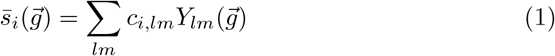

where 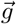 represents diffusion sensitisation gradient direction, *c_i,lm_* stand for SH coefficients and *Y_lm_* are real SH basis functions of order *m*.

#### The forward model

The acquired signal is denoted *s_jk_*, where index *j* represents an in-plane grid point within a slice denoted by index *k*. The simulation 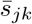 of an acquired slice *s_jk_* can be decomposed into the following steps. First, we simulate the high resolution volume 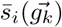 from SH coefficients *C* using the diffusion gradient 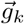 associated with the slice *k*. A point-spread-function 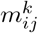 which takes into account in-plane resolution, slice thickness, position and orientation of the fetal head in the scanner space at the time of acquisition [10], is then applied to simulate the signal 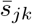 for each voxel in the slice:

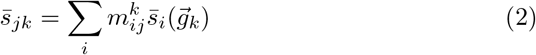

#### The objective function

The underlying SH representation *C* can then be estimated by minimising the objective function

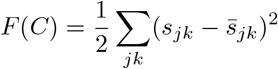

that represents sum of squared differences between acquired and simulated signals. The objective function is optimised iteratively using gradient descent with the following update equation,

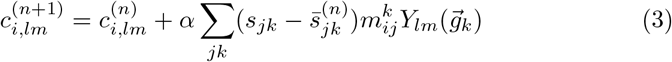

where *n* denotes the iteration number and *α* is a step size.

### 2.2. Intensity correction

Spin history effects can be modelled to a good approximation as slice-wise smoothly varying multiplicative fields. The B1 field inhomogeneity also modulates the intensities and can be considered approximatelly stationary in the scanner space and will therefore modulate data in the space of the fetal head differently as the fetus moves. The bulk for B1 inhomogeneity is corrected in the pre-processing step [19], see Fig. 1, and any residual differential multiplicative bias fields will be automatically corrected during the spin history correction. We therefore perform the intensity correction by estimating slice-wise low-frequency multiplicative fields *H* = {*h_jk_*}. In this work we use an exponential model *e^hjk^* [24, 10], though a simple multiplicative model could be used instead. The objective function then becomes

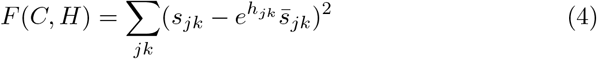

and update equation (3) will change to

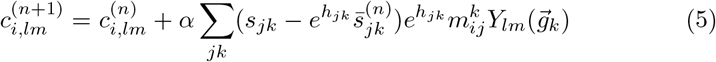

**Figure 1:**
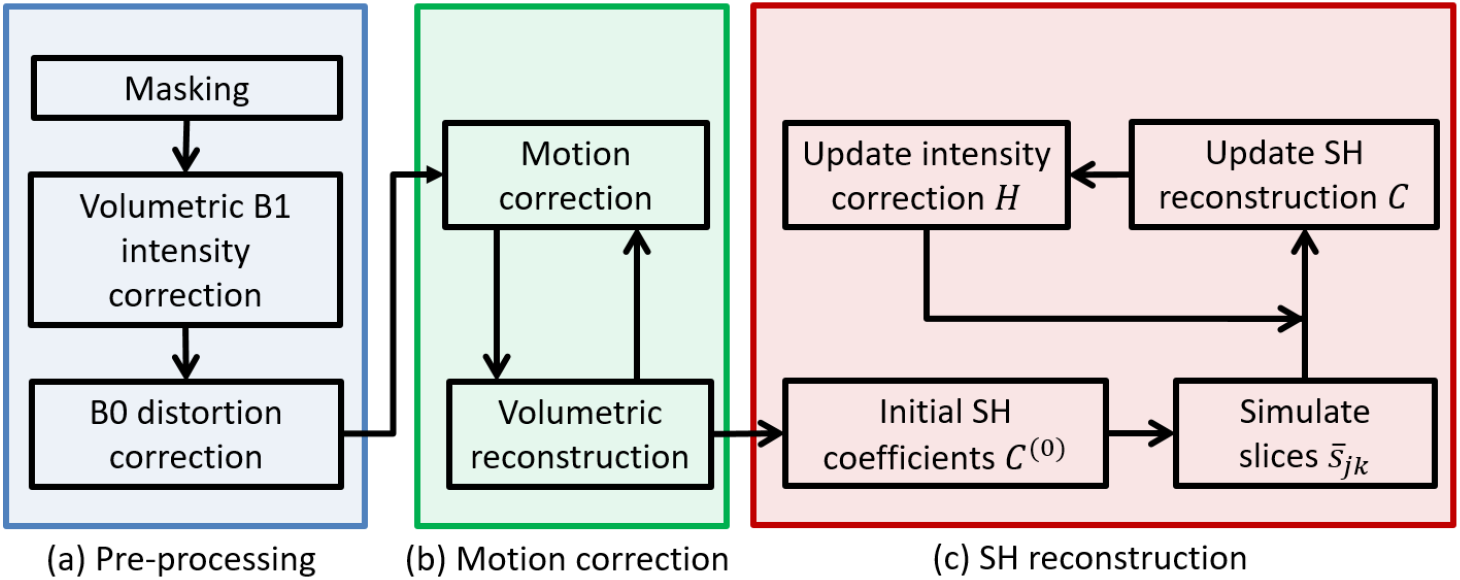
Processing pipeline

There are many ways to impose smoothness on the fields *h_jk_*, either parametric, by modelling them as a linear combination of basis functions, or non-parametric by imposing a smoothness penalty. In the case of parametric representation the smoothness is imposed by restricting the number of control points (or the basis functions), while in the non-parametric cases the covariance matrix Ψ*_k,σ_* for *h_jk_* can be defined instead. If *h_jk_* are drawn from a multidimensional Gaussian distribution *N* (0, Ψ*_k,σ_*), then these fields are modelled as Gaussian processes. This is equivalent to a parametric representation using regularised dense linear combination of Gaussian radial basis functions. We prefer the non-parametric model, because it results in more flexible space of solutions. However, it requires inversion of matrix of dimension *J_k_ × J_k_* where *J_k_* is a number of pixels in a slice *k*. Inspired by the work of Wells *et al.* [25] and proposed by us [10], we therefore approximate the estimation of the residual fields 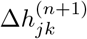 by weighted kernel regression

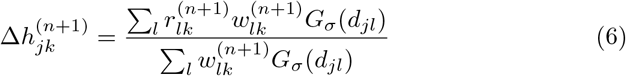

applied to residuals

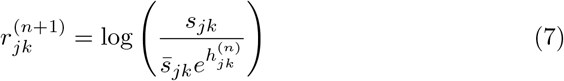

where *d_jl_* is distance between grid points *j* and *l*, and *G_σ_* represents a Gaussian kernel with standard deviation *σ* that regulates the smoothness of the intensity correction fields. The weights 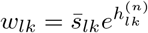 ensure that the residuals resulting from division by a small number are down-weighted as these are likely to reflect noise, while voxels with high signals drive the estimation. If voxel-wise robust statistics is used as in [10], the weights can also include posterior probability that the voxel belongs to the inlier class. The fields *h_jk_* can then be expressed as

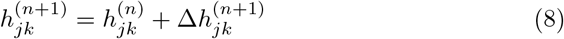

### 2.3. Distortion correction

The dMRI data are acquired as a set of diffusion sensitised and non-sensitised (*b*_0_) echo planar images (EPI). The *b*_0_ volumes have mostly T2-weighted contrast that is similar to single-shot fast spin echo (ssFSE) T2-weighted images, the modality used for structural imaging of the fetal brain which does not suffer the geometric distortion. In our previous work we have therefore proposed to register *b*_0_ stacks to ssFSE images to estimate the susceptibility induced distortion present in dMRI [15].

First, the ssFSE stacks of slices have to be motion corrected and reconstructed into a consitent structural T2-weighted volume [10]. We then register the *b*_0_ slices to the ssFSE volume by iterating between motion and distortion correction. The motion correction is performed by rigid registration of *b*_0_ slices to ssFSE volume by estimating one rigid transformation per slice. The distortion field, on the other hand, is modelled as a stationary continuous scalar 3D field, that determines the displacement of the voxels in *b*_0_ slices in phase-encoding direction. Using the estimated motion parameters, the undistorted slices are simulated from ssFSE volumes and 3D registration is then performed to aligned distorted acquired *b*_0_ slices with simulated undistorted sliced to estimate the distortion field. This is performed unsing non-rigid registration by estimating a single 3D B-spline transformation, that is further constrained to phase-encoding direction only and to obey Laplacian equation to ensure that estimated distortion field is physically plausible. The full details of the method can be found in our previous paper [15].

Based on the assumption that the distortion field can be well approximated by single 3D stationary field, the distortion field estimated from the *b*_0_ volumes can also be applied to correct the susceptibility induced distortion in the diffusion sensitised volumes.

### 2.4. The algorithm

The methodology proposed in this paper is designed for motion correction and reconstruction of single shell high angular resolution diffusion imaging (HARDI) data. Additionally, the pre-processing steps require at least one *b*_0_ volume for B1 inhomogeneity correction and motion-corrected and a reconstructed anatomical T2-weighted single shot Fast Spin Echo (ssFSE) volume as a reference for distortion correction. We also assume that B1 and B0 fields are approximately stationary throughout acquisition. The processing pipeline is summarised in Fig. 1.

#### Pre-processing

The algorithm starts by several pre-processing steps (Fig. 1a), where the dMRI is semi-automatically brain-masked by combination of thresholding and morphological operations, the B1 bias field is estimated from *b*_0_ volumes [19] and applied to correct intensities of all diffusion sensitised volumes. The stationary B0 field inhomogeneity induced distortion is estimated by registration *b*_0_ stacks to ssFSE volume of the same subject, as described in Sec. 2.3, and applied to the diffusion sensitised volumes.

#### Motion correction

The motion is estimated by coregistration of all diffusion sensitised slices irrespective of the diffusion directions using slice-to-volume registration (SVR) [10] with normalised cross-correlation as a similarity measure, as proposed previously [6]. The algorithm iterates between reconstruction of 3D volume and refinement of motion parameters (Fig. 1b).

#### Initialisation

The reconstructed high-resolution volume, that is the output of SVR [10] applied to all diffusion sensitised volumes irrespective of the sensitation gradient can be interpreted as a zero-order (*m* = 0) SH representation of dMRI signal and used for initialisation of SH reconstruction *C*^(0)^ (Fig. 1c). The estimated motion parameters are combined with acquisition specific PSF to calculate voxel specific PSFs 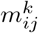 that take into account position in acquisition space and motion parameters and therefore provide a transformation between the sampled acquisition and fetal anatomical space. Spin history effect fields *h_jk_* are initialised as zero fields. Slice-specific diffusion gradients 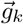 are calculated by rotating nominal volume-specific gradient vectors from the pre-defined gradient table using rotation component of the slice-specific motion parameters.

#### Reconstruction of SH representation

The isotropic SH representation *C* of the fetal dMRI signal is estimated by minimising the objective function *F* (eqn. 4) using gradient descent (GD). The algorithm proceeds by iterating between three steps:

1. Simulate slices 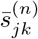 from SH representation *C*^(*n*)^ using eqn. (1) and (2).
2. Update SH representation *C*^(*n*+1)^ using eqn. (5).
3. Update intensity correction fields *H*^(*n*+1)^ using eqn. (6), (7) and (8).

If corrupted slices are present, an additional robust statistics step can be included, in a similar manner as described in our previous work for fetal structural MRI [10]. The reconstruction of SH representation is summarised in Fig. 1c.

## 3. Experiments

### 3.1. Acquisition of fetal dMRI

The methods described in Sec. 2 were aplied to three datasets with different acquisition parameters and image quality. Written informed consent was obtained from all participants and all subjects used in our experiments were healthy.

#### 1.5T anisotropic dataset

Fetal dMRI of six subjects (gestational ages 24, 26, 29, 32, 33 and 34 weeks, subjects 1-6) was acquired on a 1.5T Philips Achieva scanner using a spin echo EPI sequence with one *b*_0_ volume and 15 diffusion sensitised volumes with b = 500 s/mm^2^, TE 121ms, TR 8500ms, voxel size 2.3 × 2.3 × 3.5mm^3^, slice overlap 1.75mm. Acquisition time was approximately 5 minutes.

#### 3T anisotropic dataset

Diffusion MRI of three additional fetal subjects (gestational ages 32, 33 and 35 weeks, subjects 7-9) were acquired on a 3T Phillips Achieva scanner using a spin echo EPI sequence with three *b*_0_ volumes and 32 diffusion sensitised volumes with b = 750 s/mm^2^, TE 75ms, TR 7500ms, voxel size 2 × 2×3.5mm^3^, slice overlap 1.75mm. Acquisition time was approximately 5 minutes.

#### 3T isotropic dataset

Three dMRI fetal datasets (gestational ages 25, 30 and 37 weeks, subjects 10-12) that were acquired as a part of developing Human Connectomme Project (dHCP) during the sequence development stage were considered in this work. The datasets were acquired on a 3T Phillips Achieva scanner using spin and field echo (SAFE) EPI sequence [16] with 140 dual echo volumes on three shells (b = 0, 400 and 1000 s/mm^2^). Only the top shell and the spin echo volumes were considered in this paper, resulting in 80 b = 1000 s/mm^2^ diffusion sensitised volumes, with TE 75ms, TR 6100ms, multi-band 2, voxel size 2 × 2×2mm^3^, slice gap 0mm. Acquisition time for the whole dataset was approximately 15 minutes. For this particular dataset complex data has been denoised using random matrix theory tools [26] and a phase-based dynamic distortion correction, which takes advantage of the field echo volumes, was applied [27, 16].

### 3.2. Implementation

#### Brain masking

Brain-masking was performed using a semi-automatic approach, where all diffusion weighted volumes were registered using volumetric registration with mutual information as a similarity measure, averaged and the average image was then thresholded, followed by sequence of morphological operations to obtain the brain mask for the first stack.

#### Intensity correction

The Gaussian kernel that regulates smoothness of the bias field in the pre-processing step as well as during SH reconstruction was experimentally set to *σ* = 20mm.

#### Motion correction

We have performed three motion correction iterations, that interleaved slice-to-volume registration with reconstruction of a single volume from all dMRI stacks irrespective of the sensitation gradient. The first iteration was a constrained registration of whole stacks of slices, in the second iteration stacks were split into even and odd slices in accordance with the acquisition order and in the final iteration slices were registered to the reconstructed volume independently.

#### Reconstruction of SH representation

During SH reconstruction we performed fixed number GD iterations, that interleave simulation of the slices with updating the SH representation and intensity correction fields. Number of iterations was set to 10 for all subject to achieve a good compromise between reconstruction quality and computational time.

#### SH order

The maximum viable order for SH representation is dictated by number of volumes of different diffusion directions and the b-value. For 1.5T dataset, we have observed that the acquisition protocol and b = 500s/mm^2^ only supported a 2nd order SH model, even though theoretically, 15 directions are the minimum requirement for a 4th order model. We therefore performed 2nd order reconstructions for the 1.5T data. These are equivalent to diffusion tensor and do not support reconstruction of fiber crossings. For 3T datasets we used higher order models (4th order for 3T anisotropic and 6th order for 3T isotropic) which allow for identification of crossing fibers.

#### Running time

The algorithm was parallelised using TBB (Intel Threaded Building Blocks library) and performed on a desktop PC with 24 threads. The processing time (for motion correction and SH reconstruction with intensity correction) was approximately 15 minutes per subject.

### 3.3. Convergence of the method

We have inspected the evolution of the data consistency over iterations of SH reconstruction (Fig. 1c) as measured by the objective function (eqn. 4) and also performed visual inspection of evolving reconstructed diffusion signals and resulting fiber orientation distributions obtained by constrained spherical deconvolution [21]. These results are presented in Sec. 4.1.

### 3.4. Quantitative evaluation

We have performed the quantitative evaluation of two important elements of the pipeline: distortion correction and intensity correction for 9 fetal subjects (1.5T and 3T anisotropic datasets). Additionally, intensity correction was also evaluated for the 3T isotropic dataset. The distortion correction for this dataset was performed as a part of dHCP reconstruction pipeline and therefore not considered in this paper. For each subject we excluded 5 diffusion sensitised volumes which did not contain any corrupted slices and reconstructed the SH representation of the dMRI signal using the remaining volumes. The slices of the excluded stacks were then simulated from the reconstructed SH representation and compared to the acquired data, as proposed in previous works [10, 11].

To perform quantitative evaluation of the different element of the pipeline we calculated the root mean squared error (RMSE) between simulated and acquired stacks for three different processing pipelines: The pipeline with no distortion or intensity correction (*basic*), pipeline with distortion correction but no intensity correction (*dist*) and the full pipeline with both distortion and intensity correction (*dist+int*). To make the results more comparable for different acquisitions, the intensity ranges of all dMRI datasets were rescaled to the same average value before calculation of RMSE. The quantitative results are presented in Sec. 4.2.

### 3.5. Constrained spherical deconvolution and whole brain tractography

Evaluation of accuracy of slice to volume reconstruction of fetal MRI is difficult due to lack of ground truth. Previous works [5, 3, 8, 10, 6, 12] used simulated experiments, where motion corrupted data are simulated from non-motion-corrupted datasets and RMSE or PSNR between original and reconstructed data is calculated. However, these experiments, though valuable, cannot fully capture the challenges of reconstruction in presence of all acquisition artefacts. Additionally, a global numerical average error can be quite a poor indicator of whether the correct structure is present in the reconstructed data. As similar fetal dMRI motion correction approaches that aim at reconstruction of diffusion tensor have been already evaluated using simulated experiments in previous works [6, 12], in this paper we focus on visual assessment of the structure captured by our reconstructed fetal data. To do that, we perform constrained spherical deconvolution [21] to estimate the fiber orientation distribution (FOD) functions followed by whole brain tractography using the MRtrix3 software package [28] available from www.mrtrix.org. We assess the reconstructed data for the presence of the main white matter tracts and the 3T isotropic and anisotropic datasets are evaluated for the presence of the fiber crossings.

## 4. Results

### 4.1. Convergence of the method

Figure. 2 shows the evolution of the value of the objective function (eqn. 4) during the first 10 iterations for all 12 subjects. We can observe a decrease at every iteration and improvement slows down significantly when it reaches 10 iterations, making it a good compromise between speed and performance of the method. An example of the evolution of a fiber crossing over 10 iterations is presented in Fig. 3.

**Figure 2:**
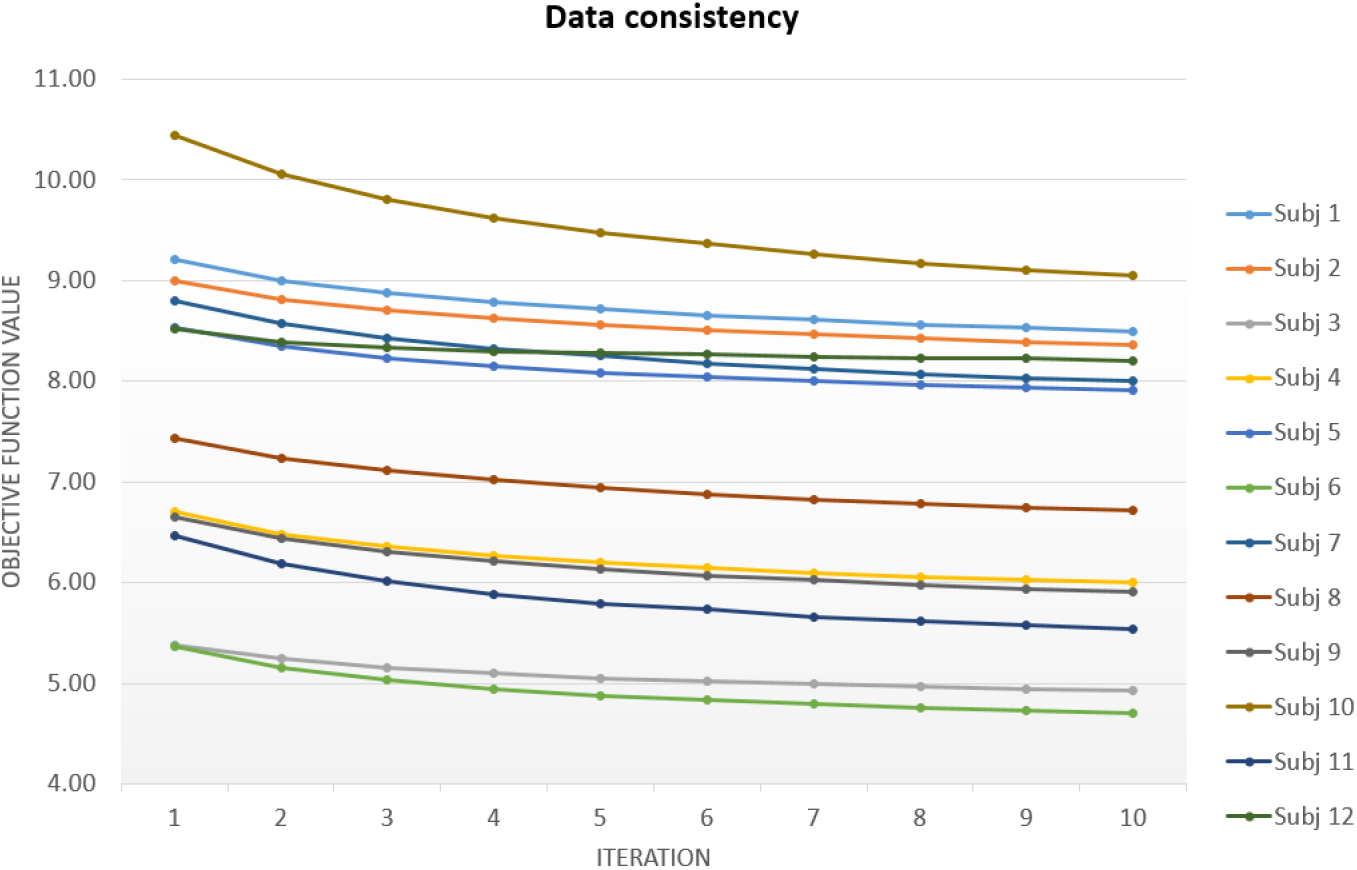
Evolution of the value of the objective fucntion during first 10 iterations for all 12 subjects.

**Figure 3:**
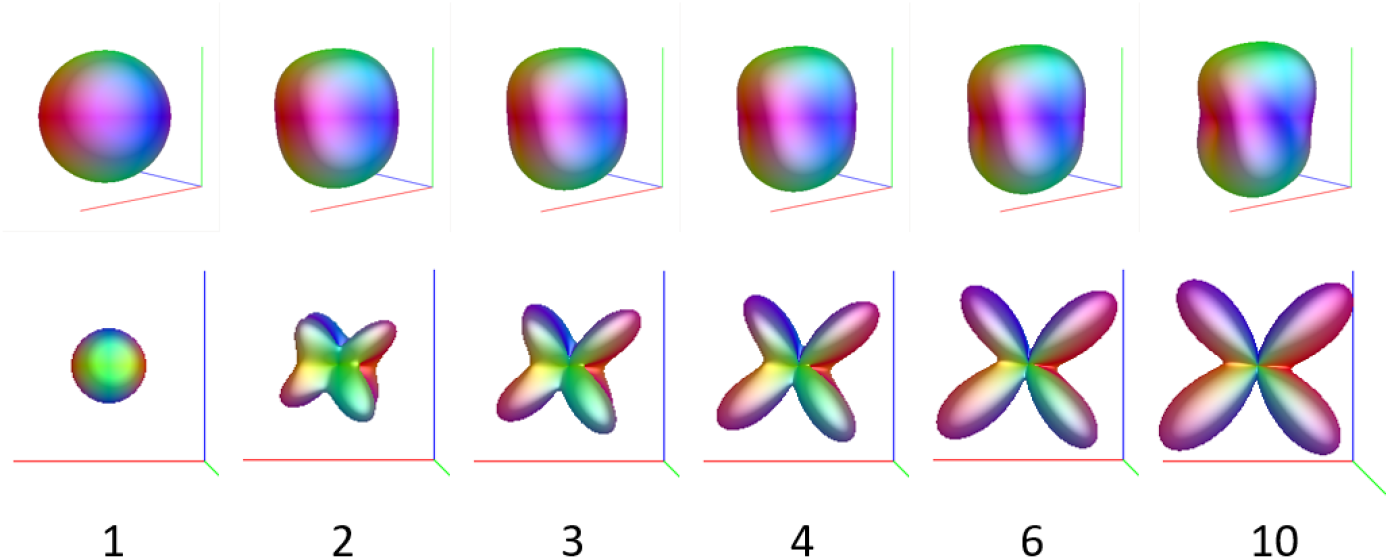
Evolution of a fiber crossing for Subject 7 depicted in Fig. 5a throughout iterations of SH reconstruction (Fig. 1c). Top row: reconstructed dMRI signal. Bottom row: Corresponding FODs obtained by constrained spherical deconvolution.

Performing further iterations might still result in small improvements in the value of objective function, however this could be caused by over-fitting to the noise in the data rather than improvements in the reconstructed SH representation. We have therefore explored evolution of the RMSE between five excluded stacks and stacks simulated from the rest of the data, as described in Sec. 3.4. We have observed that RMSE decreased for between 6-16 iterations (which varied for different subjects) and then started increasing again, indicating overfitting. Limiting the number of iteration therefore not only saves computational time but also prevents noise enhancement in the reconstructed SH representation.

### 4.2. Distortion and intensity correction

Quantitative evaluation of the effect of distortion correction and intenstity correction as described in Sec. 3.4, is presented in Tables 1 and 2. The results in Table 1 show that inclusion of distortion correction (*dist*) improves RMSE for every individual subject from the anisotropic datasets, from average value 8.21 to 8.12. A one-tailed paired student t-test confirmed that this result was significant (*p* = 1.1 × 10^−3^). Intensity correction (*dist+int*) brings substantial improvement in RMSE compared to distortion correction only (*dist*), from 8.12 to 7.30 (*p* = 9 × 10^−4^) for the anisotropic datasets, see Table 1.

**Table 1:** RMSE between acquired and simulated stacks for 9 fetal subjects from 1.5T and 3T anisotropic datasets. Pipelines compared: Basic pipeline without distortion and intensity correction (*basic*); Pipeline with distortion correction but without intensity correction (*dist*); Full pipeline with distortion and intensity correction (*dist+int*).

|  | GA | Dataset | b-value | directions | *basic* | *dist* | *dist+int* |
| --- | --- | --- | --- | --- | --- | --- | --- |
| Subj 1 | 24 | 1.5T | 500 | 15 | 9.89 | 9.79 | **9.40** |
| Subj 2 | 26 | 1.5T | 500 | 15 | 9.72 | 9.51 | **8.59** |
| Subj 3 | 29 | 1.5T | 500 | 15 | 6.19 | 6.16 | **5.49** |
| Subj 4 | 32 | 1.5T | 500 | 15 | 7.45 | 7.39 | **6.91** |
| Subj 5 | 33 | 1.5T | 500 | 15 | 9.88 | 9.77 | **8.54** |
| Subj 6 | 34 | 1.5T | 500 | 15 | 5.78 | 5.67 | **5.51** |
| Subj 7 | 32 | 3T | 750 | 32 | 8.12 | 7.94 | **6.03** |
| Subj 8 | 33 | 3T | 750 | 32 | 8.68 | 8.66 | **7.82** |
| Subj 9 | 35 | 3T | 750 | 32 | 8.20 | 8.16 | **7.09** |
| **average** |  |  |  |  | 8.21 | 8.12 | **7.30** |

**Table 2:** RMSE between acquired and simulated stacks for 3 fetal subjects from 3T isotropic dataset (3T iso). Pipelines compared: Pipeline with distortion correction but without intensity correction (*dist*); Full pipeline with distortion correction and intensity correction (*dist+int*).

|  | GA | Dataset | b-value | directions | *dist* | *dist+int* |
| --- | --- | --- | --- | --- | --- | --- |
| Subj 10 | 25 | 3T iso | 1000 | 80 | 20.23 | **11.80** |
| Subj 11 | 37 | 3T iso | 1000 | 80 | 14.61 | **8.40** |
| Subj 12 | 30 | 3T iso | 1000 | 80 | 7.69 | **6.32** |
| **average** |  |  |  |  | 14.17 | **8.84** |

The effect of intensity correction for the 3T anisotropic dataset was even more dramatic, with average RMSE for subjects 10-12 decreasing form 14.17 to 8.84, see Table 2). The RMSE without intensity correction (*dist*) is particularly high for subjects 10 and 11. This was caused by frequency drift of the scanner during lengthy acquisition resulting in intensity drift for individual diffusion volumes. Additionlly subject 10 was significantly corrupted by motion. In spite of this, the reconstruction with intensity correction (*dist+int*) resulted in good quality images, as demonstrated by the RMSE (Table 2) and visual inspection of Subject 10 (Fig. 6). For subject 12 the frequency drift problem has been resolved and the RMSE without and with intensity correction is similar to the ranges in the unisotropic datasets (Table 1).

Visual impact of the intensity correction is demonstrated in Fig. 4. The Fig. 4a show acquired stack of slices for subject 7, with inconsistencies in the brightness due to the spin history effects (shown by yellow arrows). By applying the intensity correction to the acquired data, these inconsistencies are corrected, as seen in Fig. 4b. Fig. 4c shows how intensity artefacts impact the whole brain tractography resulting in implausible and asymmetric appearance, visible in particular in coronal and transverse planes. This issue is resolved thanks to the proposed intensity correction (Fig. 4d).

**Figure 4:**
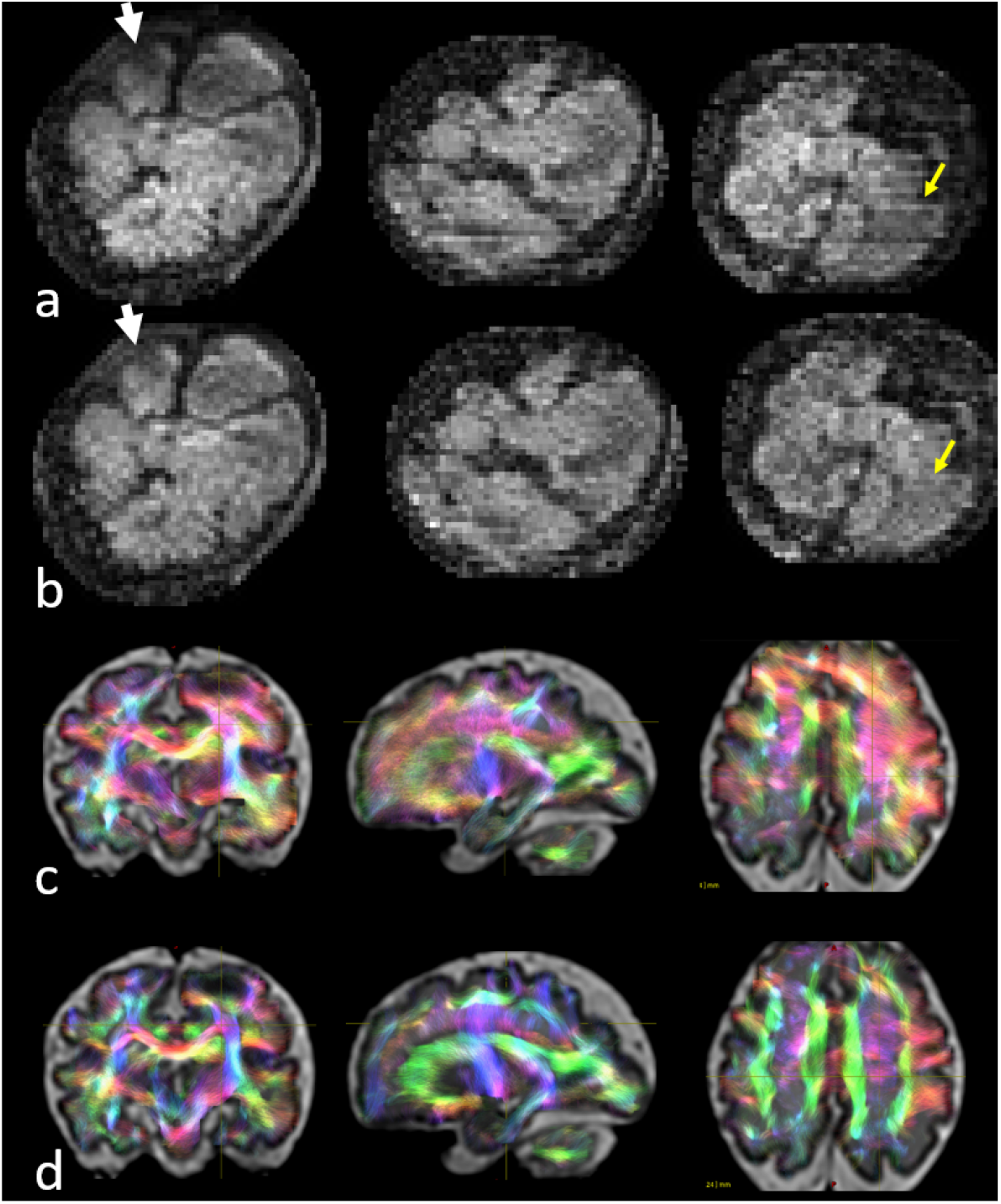
The effect of intensity correction: (a) An acquired and (b) an intensity corrected stack for Subject 7. Yellow arrows point to stripes in acquired data that has been corrected in intensity corrected data. Whole brain tractography overlayed on T2 weighted image reconstructed (c) without intensity correction (*dist*) and (d) with intensity correction (*dist-int*).

### 4.3. Assesment of reconstructed fetal dMRI

Visual inspection of the whole brain tractography confirmed that main white matter tracts could be identified in all 12 fetal subjects, including corpus callosum, cortico-spinal tract, superior longitudinal fasciculus and cingulum. The whole brain tractography of the fetal subjects 7-9 (3T anisotropic dataset) with fiber crossings of these tracts is depicted in Fig. 5.

**Figure 5:**
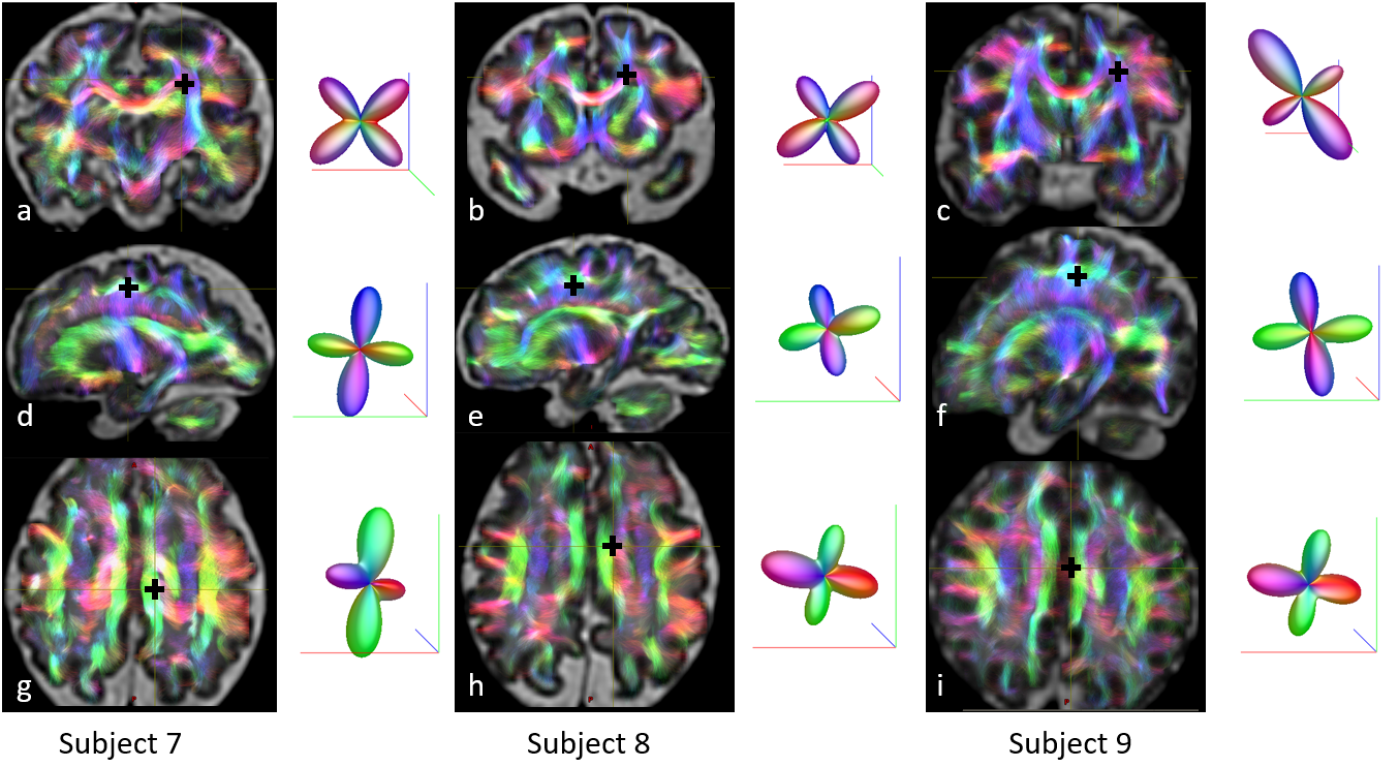
Whole brain tractography of fetal subjects 7-9 from the 3T anisotropic dataset (32, 33 and 35 weeks GA, first, second and third column respectively) overlayed on the reconstructed T2 weighted (ssFSE) anatomical image. The tractography correctly depicts main white matter tracts, colour coded as follows: Red left-right, e.g. corpus callossum (CC); Blue top-bottom e.g. cortico-spinal tract (CST); Green front-back, e.g. superior longitudinal fasciculus (SLF) and cingulum (CG). The fiber crossings are shown by black cross: First row (a,b,c): CC and CST; Second row (d,e,f): CST and SLF; third row (g,h,i): CC and CG.

In Fig. 6 we present a comparison of the fiber orientation distribution (FOD) functions for subjects 1 and 10. These subjects had similar GA at scan (24 and 25 weeks), however the acquisition protocol was very different (1.5T, 15 directions, b = 500, 2.3×2.3×3.5 mm^3^ for Subject 1 and 3T, 80 directions, b = 1000s/mm^2^, 2×2×2 mm^3^ for subject 10). In spite of significant differences of the two protocols, we can observe similar anatomy and in particular radially oriented FODs within the smooth cortical ribbon typical of young fetuses. In subject 10, however, we can also observe anatomically plausible fibre crossings (including crossings of CC and CST; and CST and SLF) thanks to our ability to fit 6th order SH model. These crossings cannot be identified for subject 1, where only 2nd order SH fit was possible.

**Figure 6:**
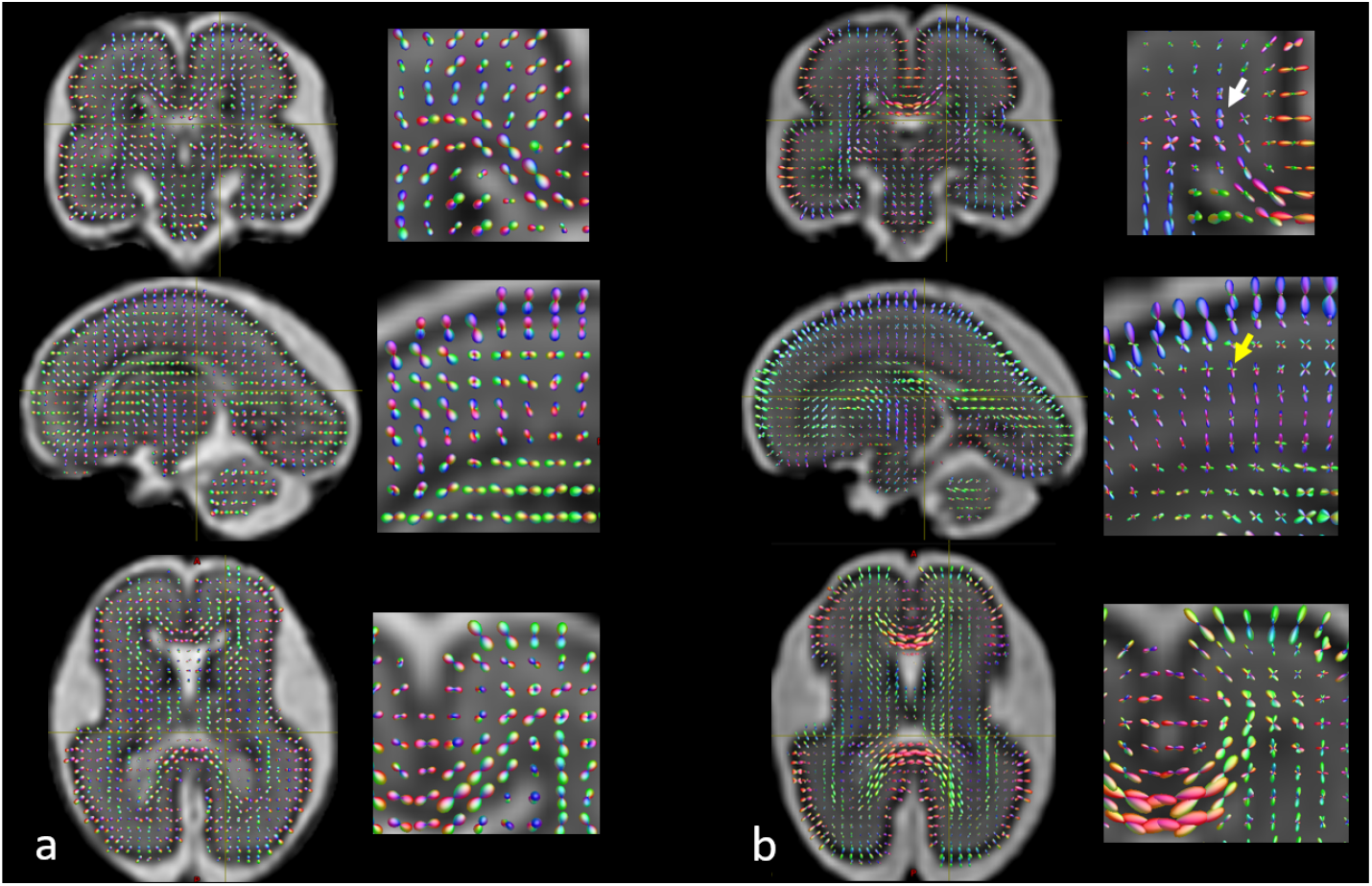
Fiber orientation distribution (FOD) functions for (a) Subject 1 (b = 500s/mm^2^, 15 directions, SH order 2, GA 24 weeks,) and (b) Subject 10 (b = 1000s/mm^2^, 80 directions, SH order 6, GA 25 weeks). We can observe radially oriented FODs within the smooth cortical ribbon typical for this stage of brain development. In subject 10, we can indentify fiber crossings, including CC and CST (white arrow) and CST and SLF (yellow arrow).

## 5. Discussion

In this paper we have proposed a novel method for reconstruction of spherical harmonics representation of diffusion MRI signal for fetal brain. We have shown that the method is applicable to datasets with various image acquisition protocols and given sufficient number of diffusion directions and b-value, we are able, for the first time, to reconstruct plausible fiber crossings in the developing fetal brain from motion corrupted fetal dMRI. We have shown that using a 4th order SH model to reconstruct dMRI with b = 750s/mm^2^ and 32 diffusion directions can already result in identification of the anatomically correct fiber crossings in the fetal brain. The SH model is flexible and can adapt to the higher number of directions and higher b-value by increasing the SH order to model higher angular resolution and we have expoited that by using 6th order spherical harmonics for reconstruction of 3T HARDI datasets.

We have also identified intensity correction as an essential feature for successful reconstruction, which sets the proposed method apart from previous works. This is partly due to the use of 3T scanners, which provide higher signal to noise ratio, making fetal scanning with b = 1000s/mm^2^ feasible, however high field also brings more significant artefacts due to inhomogeneous B0 and transmit and receive B1 field. Long T1 relaxation times in fetal brain mean that there is a trade-off between number of acquired directions and severity of spin history artefacts, thus a robust intensity correction method facilitates more efficient fetal dMRI sequences with more diffusion directions.

Our quantitative analysis has also shown that distortion correction is important for reconstruction of fetal dMRI. In this paper we have only corrected for static susceptibility induced distortion using our previously proposed method [15], though dynamic distortion correction methods [13, 16] are currently being developed. An advantage of the distortion correction approach evaluated here, however, is that it is applicable to standard dMRI fetal protocols readily available on MR scanners and does not require any additional data or specifically designed sequences.

The proposed method, in combination with high resolution fetal dMRI sequences such as those used for acquisition of dHCP fetal data, can now provide high quality consistent fetal dMRI data suitable as an input for standard analysis developed for adult brains.

## 6. Conclusion

In this paper we proposed a comprehensive framework for reconstruction of fetal dMRI using higher-order SH representation that corrects for motion, distortion, spin history and bias field artefacts. We have demonstrated that the reconstructed isotropic fetal dMRI signal can be used to perform constrained spherical deconvolution and whole brain tractography commonly used for analysis of dMRI in adult subjects. As a result, a comprehensive investigation of the in-vivo in-utero brain development is possible for the first time, including timing of the development of various white matter tracts and presence of the developmental features such as radial orientation within the fetal cortex. The higher order SH model also supports reconstruction of anatomically plausible fiber crossings in the developing fetal brain. The proposed methodology facilitates standard diffusion analysis in fetal population, opening opportunities for in depth investigation of developing fetal brain connectivity and microstructure.

## Footnotes

1 www.developingconnectome.org

## Acknowledgment

This research was funded by the ERC developing Human Connectome Project [Grant Agreement no. 319456], MRC strategic funds [MR/K006355/1] and National Institute for Health Research (NIHR) Biomedical Research Centre at Guy’s and St Thomas’ NHS Foundation Trust and King’s College London. The views expressed are those of the authors and not necessarily those of the NHS, the NIHR or the Department of Health. This work was additionally supported by the Wellcome/EPSRC Centre for Medical Engineering at King’s College London [WT 203148/Z/16/Z], the Center of Biomedical Imaging of Geneva-Lausanne Universities and EPFL, the Fondation Leenaards and Fondation Louis-Jeantet, and SNSF International Short Visit Grant (IZK0Z2 170894).

